# TPX2 expression promotes sensitivity to dasatinib in breast cancer by activating the YAP transcriptional signaling

**DOI:** 10.1101/2023.09.04.556165

**Authors:** Carlos Marugán, Beatriz Ortigosa, Natalia Sanz-Gómez, Ana Monfort-Vengut, Cristina Bertinetti, Ana Teijo, Marta González, Alicia Alonso de la Vega, María José Lallena, Gema Moreno-Bueno, Guillermo de Cárcer

## Abstract

Chromosomal instability (CIN) is a hallmark of cancer aggressiveness, providing genetic plasticity and tumor heterogeneity that allows the tumor to evolve and adapt to stress conditions. CIN is considered a cancer therapeutic biomarker because healthy cells do not exhibit CIN. Despite recent efforts to identify therapeutic strategies related to CIN, the results obtained have been very limited. CIN is characterized by a genetic signature where a collection of genes, mostly mitotic regulators, are overexpressed in CIN-positive tumors, providing aggressiveness and poor prognosis. We attempted to identify new therapeutic strategies related to CIN genes by performing a drug screen, using cells that individually express CIN-associated genes in an inducible manner. We find that the overexpression of TPX2 enhances sensitivity to the SRC inhibitor dasatinib due to activation of the YAP pathway. Furthermore, using breast cancer data from the TCGA and a cohort of cancer-derived patient samples, we find that both TPX2 expression and YAP activation are present in a significant percentage of cancer tumor samples, providing poor prognosis, being therefore putative biomarkers for dasatinib therapy.

## 1. Introduction

Chromosomal instability (CIN) is defined by a high rate of chromosome mis-segregation leading to chromosomal copy number alterations (CNAs) affecting either whole chromosomes or chromosome arm fragments [1, 2]. These CNAs ultimately result in an uneven distribution of DNA after cell division leading to aneuploidy [3]. In the context of cancer, CIN is a source of phenotypic variation that generates heterogeneity at the chromosome copy number and gene dosage level, leading to tumor progression and aggressiveness, metastasis, recurrence, and drug resistance [3–5].

CIN is rare in normal tissues but common in cancer, with 60-80% of human tumors exhibiting CIN [5, 6]. This suggests a therapeutic opportunity considering CIN as a biomarker for tumor cells, since any therapy targeting CIN would not affect healthy cells [7], and several studies evaluated the possibility of linking drug sensitivity and CIN. By screening aneuploid cells against a drug library, AICAR (AMPK activator) and 17-AAG (Hsp90 inhibitor) were identified as compounds able to specifically inhibit the proliferation of positive CIN cells [8]. Similarly, by an in silico data correlation analysis, comparing drug sensitivity for around 45000 chemicals and ploidy data from the NCI-60 database, 8-azaguanine and 2,3-diphenylbenzo[g]quinoxaline-5,10-dione (DPBQ) were identified as aneuploid-selective killing compounds [9]. Despite these initial attempts, no more advances are significant in this regard; consequently, new strategies for targeting CIN cancer cells are needed.

CIN is defined by a particular genetic signature obtained by correlating the expression of more than 10,000 genes versus a score of functional aneuploidy, as a surrogate value of CIN [10]. The top-listed genes with the highest CIN score constitute the CIN70 expression signature. Overexpression of the CIN70 signature is associated with poor clinical outcomes in several cancers. In addition, other studies identified similar CIN signatures for a variety of cancers, correlating with increased tumoral aggressiveness [11–15] and metastasis capacity [16]. The CIN signatures are particularly enriched in mitotic genes intimately linked to ensure proper chromosome segregation in each cell cycle [17, 18]. The most representative CIN70 signature gene is TPX2, an essential mitotic gene participating in the spindle assembly by activating the Aurora-A kinase [19–23]. TPX2 is overexpressed in a wide variety of tumors, and this is a hallmark of poor prognosis [17, 24, 25]. Genetic depletion of TPX2 leads to severe mitotic aberrations indicating its essentiality in cell proliferation [26]. TPX2 overexpression also leads to mitotic changes that can cause CIN and compromise cell viability [27].

The Hippo-YAP/TAZ signaling axis is an essential pathway regulating cell proliferation, cell plasticity, and organ growth during animal development. It is modulated by several inputs such as cell polarity signaling, cell–cell adhesion, cell contact inhibition, mechanotransduction, etc [28–30]. In the context of cancer, the Hippo-YAP/TAZ pathway is often deregulated, promoting adaptation and proliferation capacity to cancer cells [31, 32], and is a major sensor for adapting to elevated CIN levels [33, 34]. Indeed, elevated YAP activity correlates with CIN levels in cancer samples [35–37]. The Hippo-YAP/TAZ pathway is considered a bona fide target for cancer therapy, especially by inhibition of the YAP/TAZ transcription activity [31, 32]. In this regard, verteporfin was identified as an efficient inhibitor of the YAP/TAZ binding to the TEAD transcription factors [38], leading to an efficient downregulation of YAP/TAZ target genes. Other attempts to find YAP/TAZ inhibitors were done by screening drugs that inhibit the YAP/TAZ nuclear translocation, finding dasatinib, statins, and pazopanib in breast cancer cell lines [39, 40]. These compounds induce the proteasomal degradation of YAP/TAZ and decreased cellular proliferation.

The specific sensitivity of YAP signaling to dasatinib is based on the close interaction between YAP and the SRC family kinases (SFKs) signaling. The SFK family is comprised of eleven kinases (BLK, BRK, FGR, FYN, FRK, HCK, LCK, LYN, SRC, SRM, and YES) [41]. YAP gets its name from “YES-Associated Protein 1” because it binds to the SH3 domain of YES kinase [42]. SFKs activate YAP/TAZ either by direct YAP phosphorylation or by repressing the Hippo pathway by LATS1/2 kinases phosphorylation [43 133]. The interaction with SFKs is essential for YAP signaling activation in a wide variety of tumoral cells [43–48].

In this work, we screened MDA-MB-453 breast cancer cells that individually overexpress a collection of CIN-related genes (BIRC5, CCNB1, CCNB2, ECT2, HEC1, MAD2, PRC1, PTTG1, and TPX2) in an inducible manner by the TET-ON system, against a panel of 60 drugs already being used in the clinic or in advanced-stage clinical trials, which inhibit representative signaling pathways in cancer. We found that TPX2 overexpression provides an enhanced sensitivity towards the SRC inhibitor dasatinib, sustained on an increased YAP/TAZ signaling dependency. Moreover, after exploring the METABRIC project, and a cohort of breast cancer derived samples, we show that TPX2 expression and YAP activation are coincident biomarkers in a significant proportion of aggressive breast cancer samples, suggesting dasatinib as an alternative therapeutic avenue.

## 2. Material and Methods

### 2.1. Cell line culture

MDA-MB-453 (ATCC #HTB-131) and HEK293 (ATCC #CRL-3216) cell lines were cultured in DMEM growth medium supplemented with 10% FBS, in a humid incubator with 5% CO2 atmosphere and 37°C. Cultures were periodically tested for Mycoplasma contamination, and authenticated by the GenePrint® 10 System (Promega), and data were analyzed using GeneMapper® ID-X v1.2 software (Applied Biosystems).

### 2.2. CIN-related cDNAs cloning

The BIRC5, CCNB1, CCNB2, ECT2, HEC1, MAD2, PRC1, PTTG1, and TPX2 cDNAs were obtained from the Mammalian Gene Collection repository, amplified by the Expand High Fidelity PCR system (Roche) (primers depicted in supplementary table 2), and cloned into pENTR/D-TOPO plasmid. The final inducible expression plasmids were generated in the pLenti-CMVtight-Hygro-DEST by the Gateway LR-Clonase II Enzyme Mix system (Invitrogen). All cDNA amplification and cloning steps were validated by DNA sequencing.

### 2.3. Lentiviral particle production and generation of the TET-ON inducible expression system cell line

HEK293 cells were transfected with a mixture of the 3rd generation lentiviral packaging plasmids containing: 2 μg of Rev (pRSV-Rev, Addgene #12253), 2 μg of Gag and Pol (pMDLg/pRRE, Addgene #12251), and 2 μg of VSV-G envelope expressing plasmid (pMD2.G, Addgene #12259), and either 2 μg of the Tet-On-3G transactivator (pLVX Tet3G, Clontech) or 2 μg of the pLenti-CMVtight-Hygro-DEST plasmid with each cDNA of interest, and 20 μl Lipofectamine 2000 (Invitrogen) in Opti-MEM (Gibco). Viral supernatants were retrieved after 24, 48, and 72 hours, cleared through a 0.45 μm pore-size filter and stored at -80°C.

MDA-MB-453 cells were firstly transduced with the pLVX Tet3G viral particle for the rtTA transactivator expression, and selected in 400 μg/ml of geneticin (Gibco). rtTA-expressing cells were then infected with the pLenti-CMVtight DEST plasmids expressing each cDNA of interest, and selected in 50 μg/ml of Hygromycin B. Cell stocks were amplified and stored in liquid nitrogen. MDA-MB-453 Tet-ON cells were grown in the presence of 0.01 μg/ml, 0.1 μg/ml, or 1.0 μg/ml of doxycycline (DOX) to test each cDNA expression.

### 2.4. Standard of care drug library screening

A total of 60 kinase inhibitors, including staurosporine as a positive control, were used for the drug screen experiments. These 60 inhibitors represent a collection of standards of care for many different signaling pathways and cellular processes (supplementary table 1). The MDA-MB-453 Tet-ON cell lines were dispensed in two 384-well Poly-D-lysine Biocoat plates, per cDNA of interest. For cDNA induction, DOX was added to one replicate plate at a final concentration of 0.1μg/ml, and plain growth media to the other replicate plate. After overnight incubation, 10 serial dilutions of the compound library (from 20 μM to 0.001 μM) were added to each plate, and cells were then allowed to grow for two population-doubling times. DOX and drug treatment was renewed every 48 hours.

After gene induction and drug incubation, cells were fixed in 70% cold ethanol and stained with 0.4μg/ml of DAPI. The number of cells in each well was counted with Acumen eX3 (TTP LabTech), normalized versus the DMSO control wells, and then the ratio noDOX/+DOX was calculated to evaluate the effect of each drug. Prism (GraphPad) was used to generate and analyze the scatter plots. IC50s were calculated using Genedata Screener software using the normalized cell number in each well.

### 2.5. Cell colony formation and drug treatments

MDA-MB-453 TET-ON/TPX2 cells were seeded in triplicates in 12 well plates. TPX2 expression was induced by adding 0.1μg/ml or 0.01μg/ml of DOX. The next day plates were treated with DMSO or the mentioned inhibitor concentration. DOX and inhibitors treatment were renewed every 2 to 3 days. After 12 to 14 days, the plates were fixed in 4% formaldehyde in PBS, stained with Giemsa and scanned in an Epson V800 scanner. The colony area was quantified using Image J (NIH) and its ColonyArea plugin [49].

### 2.6. Protein extraction and Immunoblotting assays

Cells were lysed in RIPA buffer (37 mM NaCl, 0.5% NP-40, 0.1% SDS, 1% Triton X-100, 20 mM Tris-HCl, pH 7.4, 2 mM EDTA, 10% glycerol, supplemented with protease and phosphatase inhibitory cocktails (SIGMA)) on ice during 20 minutes, and protein lysates clarified by 30-minute centrifugation at 14,000 rpm. Protein concentration was quantified using Pierce™ BCA Protein Assay kit (Thermo Fisher Scientific). Proteins were separated in Novex^TM^ 4%-20% tris-glycine acrylamide gels (Invitrogen) and transferred to nitrocellulose membranes (BioRad). Blotted proteins were blocked in 5% non-fat milk in PBS-T (PBS with 0.05% Tween-20), and probed with the corresponding primary antibody (supplementary table 3). Secondary antibodies coupled to fluorescent IRDye680 were incubated for 45 minutes, and scanned with the Odyssey Infrared Imaging System (Li-Cor Biotechnology).

### 2.7. Flow cytometry cell cycle analysis

Cells were trypsinized, fixed with cold 70% ethanol, and resuspended in PBS-T (PBS + 0.03% TritonX-100). DNA was stained with 1 μg/ml DAPI and DNA profile data was retrieved using a FACSCantoII device and analyzed using FACSDiva software (Becton Dickinson).

### 2.8. Cell Immunofluorescence and YAP signal quantification

Cells were fixed in 4% methanol-free formaldehyde (PolySciences) in PBS, permeabilized with cold methanol, blocked with 10% fetal bovine serum in PBS-T (PBS + 0.03% TritonX-100), and incubated with primary antibodies against phospho-histone H3-Ser10 (Cell Signaling Technology #3377s), total YAP (Santa Cruz Biotechnology #sc-271134), or active-YAP (Abcam #ab205270) diluted in PBS-T. A secondary antibody coupled to the Alexa488 dye (Invitrogen – Molecular Probes) was used. DNA was counterstained with 0.1 μg/ml DAPI, and cells were finally mounted in glass slides using ProLong Diamond antifade mounting media (Thermo Fischer). Pictures were obtained using a NIKON 90-eclipse microscope. YAP nuclear/cytoplasm ratio was calculated using the ImageJ Intensity Ratio Nuclei Cytoplasm tool (<u>RRID:SCR_018573</u>)

### 2.9. Quantitative Real-Time PCR

Single-gene qRT-PCR analysis was done using FAM-MGB TaqMan probes (Invitrogen) specific for Cyr61 (Hs00155479_m1), CTGF (Hs01026927_g1), and TEAD4 (Hs01125032_m1) genes. RNA was extracted with Trizol and column-purified with the Absolutely RNA miniprep kit (Stratagene). Both cDNA synthesis and PCR amplification were done with the SuperScript III one-step RT-PCR system (Invitrogen). TaqMan probes for the housekeeping genes ACTB (Hs01060665_g1) or HPRT1 (Hs02800695_m1) were used as normalization controls.

### 2.10. METABRIC in silico data analysis

Clinical and expression data of a cohort of 1536 patients were downloaded from the METABRIC breast cancer project [50]. Patients were classified according to TPX2 expression with a membership probability estimated by Bootstrap [51]. Gene Set Enrichment Analysis (GSEA) was performed by comparing high versus low TPX2 expressing samples. The top 25 terms of C6: Oncogenic Signature were analyzed. Additionally, we performed a GSEA with the TAZ/YAP signature [52]. FDR q< 0.05 was considered significant. Samples classified according to TPX2 expression were also classified with the YAP/TAZ-signature expression as in [53]. Briefly, after quantile normalization, a z-score was calculated for the expression data. Genes included in the YAP/TAZ signature were extracted, and each sample’s combined score was calculated as the sum of the individual expression z-scores. Samples were then classified as having a low TAZ/YAP signature if the combined score was less than the combined mean, and as having a high TAZ/YAP signature if the combined score was greater than the combined mean. Contingency tables were analyzed by a Fisher’s Exact Test.

### 2.11. Human Breast Cancer tumoral samples IHC analysis

A total of 99 paraffin-embedded grade 3 Invasive Ductal Breast Carcinoma (IDBC), molecularly classified according to the NCCN breast carcinoma guidelines, were subjected to immunohistochemistry (IHC) analysis. Samples were obtained from the MD Anderson Foundation Biobank (record number B.0000745, ISCIII National Biobank Record-Madrid) between 2003 and 2014. The mean age of the patients was 59.6 ± 9.1 years. Non-tumoral breast tissue was analyzed as an internal control. The study was approved by the local ethical committee from each institution, and complete written informed consent was obtained from all patients. Written informed consent was obtained from all patients before enrollment. The collected tumor samples were fixed in 4% formaldehyde and embedded in paraffin. Immunohistochemistry was done using TPX2 antibody (Abcam ab32795), phosphor-Ser127-YAP (Abcam ab76252), and total YAP (Santa Cruz Biotechnology sc-271134). Briefly, tumor sections were deparaffinized using the BOND Dewax Solution (Leica Biosystems) and rehydrated with serial passage through changes of decreasing ethanol solutions. Immunostaining was performed on a BOND RXm autostainer (Leica Biosystems) using BOND Epitope Retrieval Solutions (Leica Biosystems) and the BOND Polymer Refine Detection kit (Leica Biosystems), following the manufacturer’s instructions. Stained slides were counterstained with hematoxylin, scanned with an Aperio CS2 scanner (Leica Biosystems), and analyzed with the Aperio eSlide Manager software (Leica Biosystems). A semiquantitative scoring of each marker based on staining intensity and the percentage of stained tumor cells was applied by two independent observers blinded to clinical data.

### 2.12. Statistical analysis and DepMap in silico data browsing

Statistical analysis was performed using Prism software (GraphPad). The statistical significance was evaluated using either one-way ANOVA, two-way ANOVA, or correlation and linear regression. Data were plotted as mean ± SD or mean ± SEM. Probabilities of less than 0.05 were considered statistically significant: p < 0.05 (*); p < 0.01 (**); p < 0.001 (***); p < 0.0001 (****).

For tumor samples, statistical analysis was performed using SPSS Statistics 28.0.1.1 (SPSS Inc., Chicago, IL). The chi-square or Fisher’s exact tests were used to test associations between categorical variables. All tests were two-tailed, and 95% confidence intervals (CIs) were used. Values of p < 0.05 were considered statistically significant.

DepMap portal (www.depmap.org) [54] browsing was used to correlate mRNA expression data in a cohort of breast cancer cell lines, expressed in Transcript Per Million (TPM), of TPX2 versus SFK kinases, YAP/TAZ related genes, proliferative or CIN associated genes. Similarly, drug response data to dasatinib was retrieved in the format of Area Under the Curve (AUC), and correlated to mRNA expression data or the aneuploidy score as a CIN surrogate.

## 3. Results

### 3.1. Drug screening in TPX2-inducible expressing MDA-MB-453 cells identifies dasatinib as a selective hit

We performed a drug screening for mitotic and CIN-related genes by creating a TET-ON inducible gene expression system in the breast cancer cell line MDA-MB-453. Individual expression of nine different mitotic cDNAs (BIRC5, CCNB1, CCNB2, ECT2, HEC1, MAD2, PRC1, PTTG1, and TPX2) was verified by western blot after doxycycline (DOX) addition (supplementary figure 1A). The validated expressing cells were then subjected to the drug sensitivity screening using a drug panel of 60 small compounds (supplementary table 1), in the absence (no overexpression of cDNA) or presence of DOX (overexpression of cDNA). Hence, we compare in the same genetic background, the differences in drug response upon overexpression of a particular gene of interest. The drug response data, after treatment with 10 concentration dilutions of each tested compound, was obtained by establishing a noDOX/+DOX proliferation ratio (supplementary figure 1B) and plotted to identify values above 1.5 indicating that +DOX induced a 50% decrease in cell proliferation. Out of the nine different cDNAs, inducible expression of TPX2 (Figure 1A) provided a strong differential response towards the SRC kinase inhibitor dasatinib (Figure 1B). This dasatinib response is also observed by retrieving data from the DepMap portal, correlating the TPX2 expression and the response to dasatinib in a cohort of breast cancer cell lines (Figure 1C). Similar correlation data was obtained in breast and lung cancer cell lines from the Cancer Therapeutics Response Portal from the Broad Institute [55] (supplementary figure 1C).

**Figure 1.**
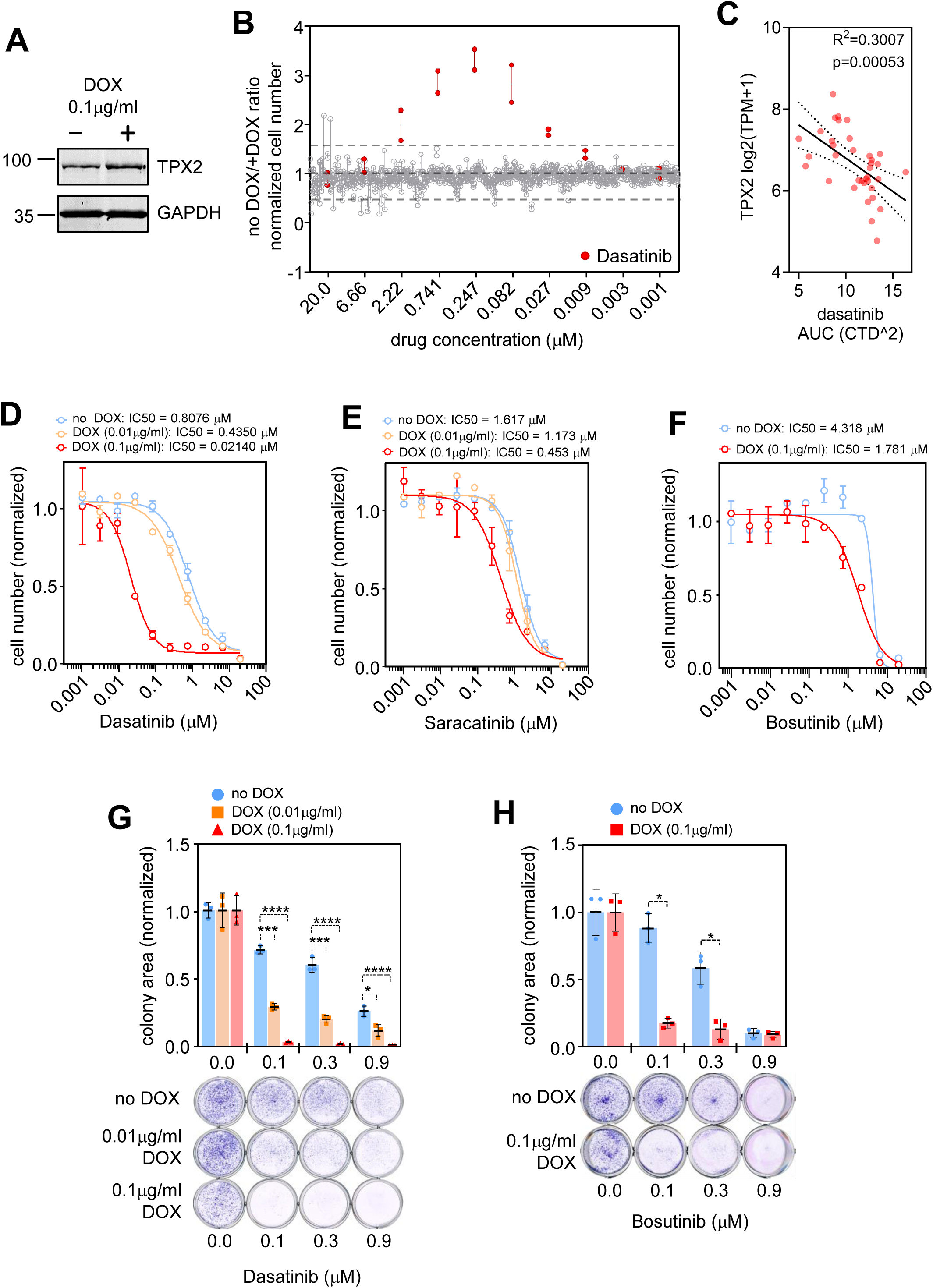
TPX2 inducible overexpression leads to increased sensitivity to SRC kinase inhibitors. **A).** MDA-MB-453 cells increase TPX2 expression upon the inducible activation of the TET-ON system by doxycycline (DOX). **B).** Drug screening scatter plot of noDOX / +DOX ratios showing the response at each drug concentration tested in MDA-MB-453 cells. Ratios for each compound (duplicates) are represented as grey dots, except dasatinib (highlighted in red). **C).** DepMap portal retrieved data, from breast cancer cell lines, showing the correlation between TPX2 expression (transcript per million – TPM) vs the sensitivity to dasatinib (area under the curve - AUC). The lower AUC, the more sensitivity to the drug. **D).** 10-point Concentration-Response Curves (CRC) and IC50 calculation of dasatinib in MDA-MB-453 cells, expressing TPX2, during 3 doubling times. Mean cell number normalized vs. DMSO +/- SEM (blue = noDOX control, orange = 0.01 μg/ml DOX, red = 0.1 μg/ml DOX).**E).** 10-point CRC and IC50 calculation of saracatinib in MDA-MB-453 cells, expressing TPX2, during 3 doubling times. Mean cell number normalized vs. DMSO +/- SEM (blue = noDOX control, orange = 0.01 μg/ml DOX, red = 0.1 μg/ml DOX). **F).** 10-point CRC and IC50 calculation of bosutinib in MDA-MB-453 cells, expressing TPX2, during 3 doubling times. Mean cell number normalized vs. DMSO +/- SEM (blue = noDOX control, red = 0.1 μg/ml DOX). **G).** Colony formation assay, during two weeks, in MDA-MB-453 cells upon 0.1μM, 0.3μM, and 0.9μM of Dasatinib treatment. The colony area is normalized vs. the DMSO-treated cells +/- SD. Two-way ANOVA with Tuckey multiple comparisons test: p<0.0001 (****), p<0.001 (***), p<0.01 (**), p<0.05 (*). (blue = noDOX control, orange = 0.01 μg/ml DOX, red = 0.1 μg/ml DOX). **H).** Colony formation assay, during two weeks, in MDA-MB-453 cells upon 0.1μM, 0.3μM, and 0.9μM bosutinib treatment. The colony area is normalized vs. the DMSO-treated cells +/- SD. Two-way ANOVA with Tuckey multiple comparisons test: p<0.05 (*). (blue = noDOX control, red = 0.1 μg/ml DOX).

To further validate this data, we performed Concentration-Response Curves (CRC) to calculate the IC50 drug response of dasatinib (Figure 1D) and other SRC kinase inhibitors such as saracatinib (Figure 1E) and bosutinib (Figure 1F). TPX2 expression provides enhanced sensitivity to the three tested drugs in a dose-dependent manner. Additionally, we tested the response to SRC inhibitors by performing cell colony formation assays in the presence of dasatinib and bosutinib (Figure 1G, H), again demonstrating that TPX2 expression increases the cellular sensitivity to SRC inhibitors. Interestingly, TPX2 expression does not provide any differential response to imatinib (supplementary figure 2A), a specific inhibitor of the BRC-ABL translocation, which has very limited activity towards SRC kinase [55, 56], indicating that the mechanism associated with TPX2 expression might exclusively rely on the SFK kinase family and not on the BRC-ABL translocation.

Since TPX2 expression is also a bona fide marker for CIN [10, 57, 58], we correlated dasatinib sensitivity versus the aneuploidy score (as a surrogate indicator for CIN) using the breast cancer cell dataset in the DepMap portal. TPX2 sensitivity does not correlate with aneuploidy or other CIN-related genes such as PRC1 and FOXM1 (supplementary figure 2B). Similarly, as TPX2 is also considered a proliferative gene [58, 59], we correlated dasatinib AUC data versus the expression of genes closely related to proliferation such as MKI67, PCNA, and MCM2 (supplementary figure 2C). Only MKI67 shows a correlation trend, but not significant, in the breast cancer cell line cohort. PCNA and MCM2 show no correlation with dasatinib AUC indicating that cell proliferation index is not a determinant for dasatinib inhibitory effect.

### 3.2. SRC kinase signaling pathway evaluation upon TPX2 induction

To understand the increased sensitivity to dasatinib in TPX2-expressing breast cancer cells, we first checked if the expression of SFK kinases correlates with TPX2 expression. TPX2 has a strong expression correlation with SRC, YES, FYN, ABL1, and ABL2 (Figure 2A), which are SFKs known to exert a prominent role in breast cancer cell lines [60]. Off note, TPX2 and LCK expression do not correlate, most probably because there is very low LCK expression in breast cancer cell lines, which is concomitant with its lymphocyte specificity expression [61].

**Figure 2.**
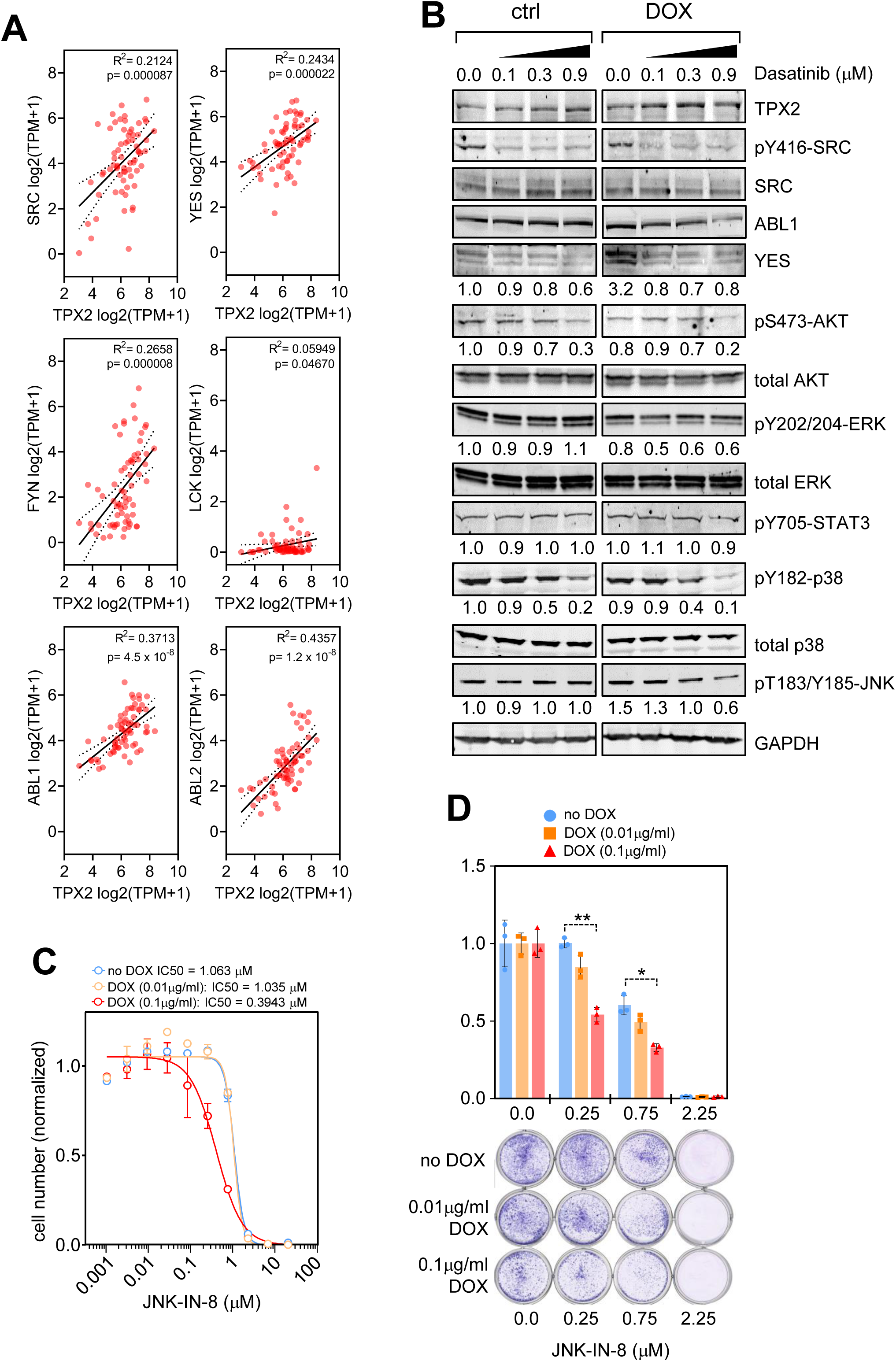
Evaluation of SFK signaling upon TPX2 inducible overexpression. **A).** Correlation analysis of TPX2 expression levels of and SFK family kinases (SRC, YES, FYN, LCK, ABL1, and ABL2), using breast cancer cell lines data retrieved from the DepMap portal. **B).** SRC signaling analysis by western blot of MDA-MB-453 cells expressing TPX2 (DOX) or control cells (ctrl), and treated with 0.1μM, 0.3μM, and 0.9μM of Dasatinib. Numbers at the bottom of the WB panels represent normalized band intensity quantified with the LiCor Odyssey 2.0 software. GAPDH expression levels are used as a loading control. **C).** 10-point CRC and IC50 calculation of the JNK kinase inhibitor JNK-IN-8 in MDA-MB-453 cells, expressing TPX2, during 3 doubling times. Mean cell number normalized vs. DMSO +/- SEM (blue = noDOX control, orange = 0.01 μg/ml DOX, red = 0.1 μg/ml DOX). **D).** Colony formation assay, during two weeks, in MDA-MB-453 cells upon 0.1μM, 0.3μM and 0.9μM of JNK-IN-8 treatment. The colony area is normalized vs. the DMSO-treated cells +/- SD. Two-way ANOVA with Tuckey multiple comparisons test: p<0.01 (**), p<0.05 (*). (blue = noDOX control, orange = 0.01 μg/ml DOX, red = 0.1 μg/ml DOX).

We then tested the expression and activation levels of SFKs (Figure 2B). There are no significant changes in the SRC or ABL1 protein levels upon TPX2 expression, but a strong increase in YES kinase levels that drops down upon dasatinib addition. Yet, we do not detect any activation of SFK signaling by testing the pTyr416 residue levels [62].

We also tested some major SRC downstream signaling pathways such as PI3K/AKT [63–65] and RAS/MEK/ERK [66–68] (Figure 2B). Despite there being a description of a TPX2 and PI3K axis connection [69], we do not observe a significant alteration in PI3K signaling, since dasatinib treatment reduces AKT pSer473 levels equally in control and TPX2-expressing cells. Concomitantly, when TPX2-expressing cells are subjected to inhibition of PI3K (BYL-719) or AKT (AZD-5363), there are no changes in cell growth (supplementary figure 3A, B). Regarding RAS/MEK/ERK signaling, TPX2 expression leads to a slight reduction in ERK signaling activation (pTyr202/Thr204) that is accentuated upon dasatinib treatment, whereas control cells do not show any phosho-ERK modification (Figure 2B). There is also a differential response to RAS/MEK/ERK inhibitors such as dabrafenib (BRAF) or trametinib (MEK) (supplementary figure 3C, D). A possible explanation for this increased response to RAS/MEK/ERK inhibitors is that CIN cells rely on the RAS/MEK/ERK axis to cope with the aneuploidy-induced cellular stress [70]. Moreover, the increased sensitivity to RAS/MEK/ERK inhibition is not as strong as with dasatinib, probably indicating that the RAS/MEK/ERK axis is not the unique modification for dasatinib-acquired sensitivity.

SRC kinases also activate the STAT signaling pathway [71, 72], but we do not detect any alteration in STAT3-pTyr705 phosphorylation upon TPX2 expression (Figure 2B). The stress signaling kinases p38 and JNK are also known downstream effectors of SRC activity [73–76]. We do not observe any alteration in p38 activation (pTyr182) upon TPX2 induction, and dasatinib suppresses p38 activation equally in control and TPX2-expressing cells. On the contrary, we observe a differential response in the activation of JNK kinase (Figure 2B). TPX2 induction leads to increased activation phosphorylation at JNK-Thr183/Tyr185, and more importantly, this activation is reduced upon dasatinib addition only in the TPX2-expressing cell. To further verify the impact of JNK in the TPX2-mediated signaling, we tested cell survival upon JNK inhibition (Figure 2C, D), and observed that TPX2-expressing cells are more sensitive to the inhibitor JNK-IN-8 than control cells, indicating that JNK might participate in the dasatinib-acquired sensitivity upon TPX2 expression.

### 3.3. TPX2 expression leads to elevated YAP/TAZ signaling activity that is reverted by dasatinib addition

SFK kinases are important drivers of YAP/TAZ activity in several cancer types leading to tumor growth and metastasis [46, 77], and dasatinib efficiently inhibits the YAP/TAZ transcriptional survival signaling [39, 40, 45, 46, 78, 79]. Moreover, a proposed mechanism for YAP activation by SRC is mediated by the stress kinase JNK [45]. Since we observed an increase in YES kinase expression and differential activation of JNK upon TPX2 expression, we focused on the possibility of a YAP/TAZ signaling modulation upon TPX2 expression. We confirmed that YAP and TAZ expression strongly correlates with TPX2 expression in breast cancer cell lines (Figure 3A).

**Figure 3.**
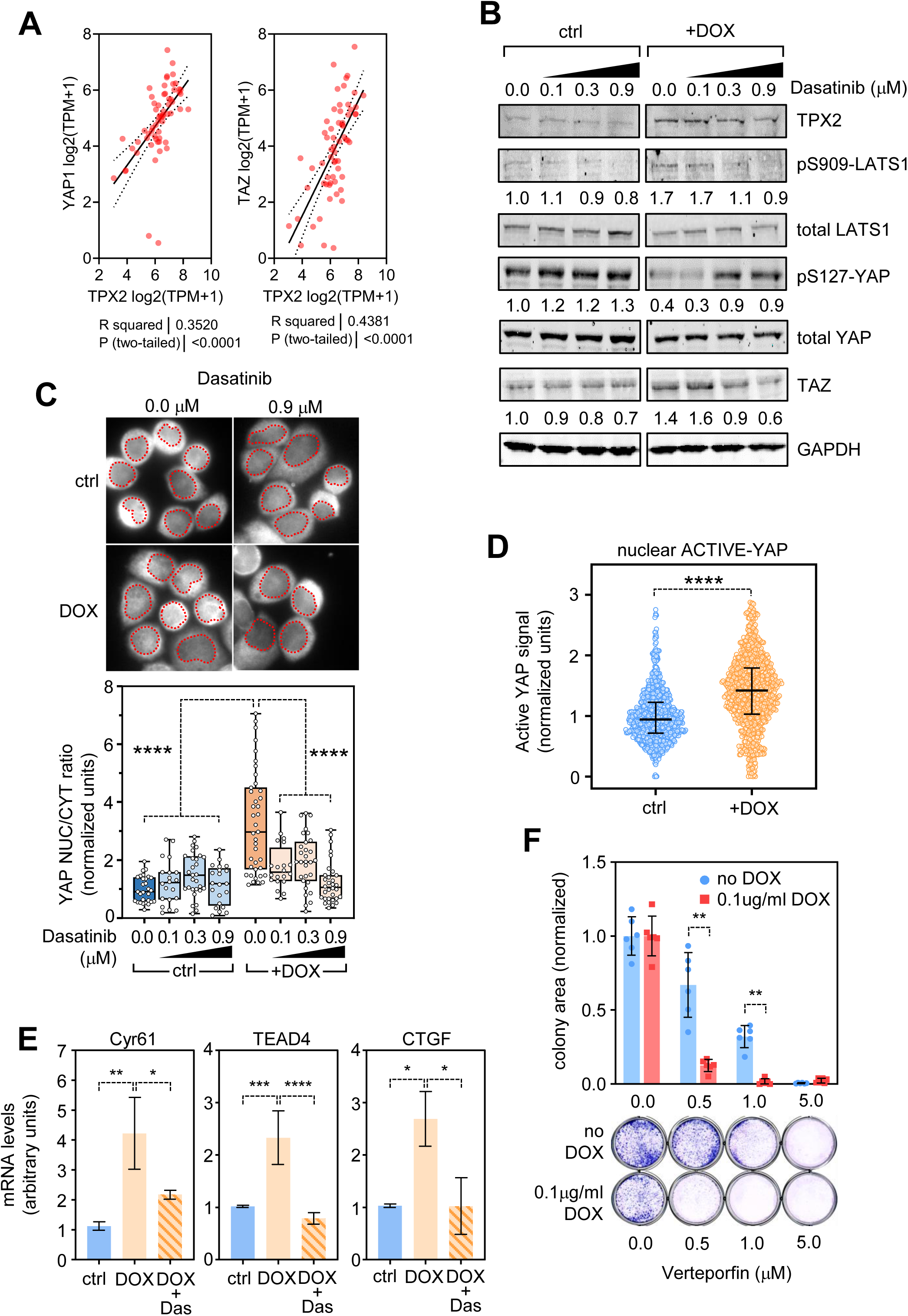
TPX2 expression leads to increased YAP signaling. **A).** Correlation analysis of TPX2 expression levels of and the YAP1 and TAZ transcription factors, using breast cancer cell lines data retrieved from the DepMap portal. **B).** Hippo/YAP signaling analysis by western blot of MDA-MB-453 cells expressing TPX2 (DOX) or control cells (ctrl), treated with 0.1μM, 0.3μM, and 0.9μM of Dasatinib. Numbers at the bottom of the WB panels represent normalized band intensity quantified with the LiCor Odyssey 2.0 software. GAPDH expression levels are used as a loading control. **C).** YAP nuclear/cytoplasm ratio analysis in MDA-MB-453 cells upon TPX2 overexpression and dasatinib treatment. Normalized data to the control untreated cells is represented in the histogram showing control cells (blue bars - ctrl) or TPX2 expressing cells (orange bars - DOX) at 0.1μM, 0.3μM and 0.9μM of Dasatinib (light colored bars). Each dot represents a single microscopy field. One-way ANOVA test comparisons: p<0.001 (****) **D).** Immunofluorescence of active-YAP in MDA-MB-453 cells upon TPX2 expression with DOX. The nuclear signal is quantified with ImageJ software, and normalized vs. control data is plotted. Unpaired T-Test analysis: p<0.0001 (****). **E).** RT-qPCR gene expression test of Cyr61, CGTF, and TEAD4 as surrogate markers of YAP transcription activity, showing mean mRNA levels (arbitrary units) +/- SD. One-way ANOVA with Tukey’s multiple comparisons post hoc test *p < 0.05; **p < 0.01, ***p < 0.001. **F).** Colony formation assay, during two weeks, in MDA-MB-453 cells upon 0.5μM, 1.0μM, and 5.0μM of verteporfin treatment. The colony area is normalized vs. the DMSO-treated cells +/- SD. Two-way ANOVA with Tuckey multiple comparisons test: p<0.01 (**). (blue = noDOX control, red = 0.1 μg/ml DOX).

TPX2 expression leads to a reduction in the YAP inhibitory phosphorylation at Ser127, elevated levels of TAZ, and increased pSer909-LATS1 (Figure 3B), which is in accordance with the described YAP-LATS1 feedback loop [80], and with previous data showing that TPX2 also leads to YAP/TAZ stabilization and activation [81, 82]. This YAP activation is reversed when dasatinib is added to TPX2-expressing cells, restoring YAP-Ser127 phosphorylation and recovering basal levels of TAZ protein (Figure 3B). Concomitantly, TPX2 induction promotes YAP nuclear shuttling in MDA-MB-453 cells, which is reversed by dasatinib (Figure 3C). This data coincides with elevated levels of active-YAP signal in the nucleus upon TPX2 induction (Figure 3D). We confirmed the TPX2-dependent YAP/TAZ activation with increased expression of the surrogate genes Cyr61, TEAD4, and CTGF and its suppression by the addition of dasatinib (Figure 3E). The DepMap portal also shows that TPX2 expression levels in breast cancer cell lines strongly correlate with the expression of YAP/TAZ surrogate genes such as CTGF, Cyr61, ANKRD1, or CRIM1 (supplementary figure 4A). Finally, the YAP signaling inhibitor verteporfin [38] significantly inhibits the growth of TPX2-expressing cells compared to control cells, demonstrating that TPX2 expression leads to a YAP-dependent growth (Figure 3F).

In summary, our data show that TPX2 leads to activation of the YAP/TAZ survival pathway, which confers enhanced sensitivity to dasatinib.

### 3.4. Dasatinib leads to a mitotic arrest in TPX2-expressing cells

Changes in TPX2 expression, either down or upregulation, lead to mitotic aberrations that prevent proper chromosome segregation, leading to CIN [26, 27]. On the other hand, although SFKs allow progression through the early phases of the cell cycle [23] and dasatinib efficiently arrests the cell cycle in G0/G1 [83–86], in recent years there is strong evidence that SFKs are also functional during mitosis [67, 87–91].

We thus evaluated the cell cycle status upon TPX2 expression and dasatinib intervention in MDA-MB-453 cells. As expected, dasatinib addition to non-TPX2 expressing cells leads to a significant reduction in the G2/M proportions, whereas TPX2 induction leads to a G2/M cell cycle arrest (Figure 4A). Interestingly, when TPX2-expressing cells are treated with dasatinib, there is a strong enhancement of the G2/M phase arrest in a dasatinib and TPX2 dose-dependent manner (Figure 4A). This cell cycle arrest is mostly due to a mitotic arrest, as depicted by pSer10-H3 staining and quantification (Figure 4B, C). The mitotic arrest led by dasatinib is corroborated by increased expression of mitotic proteins such as Cyclin B (CycB) or Aurora kinase A (AurKA), and also a notable increase in phospho-AurKA signal at the activatory residue Thr288 (Figure 4C).

**Figure 4.**
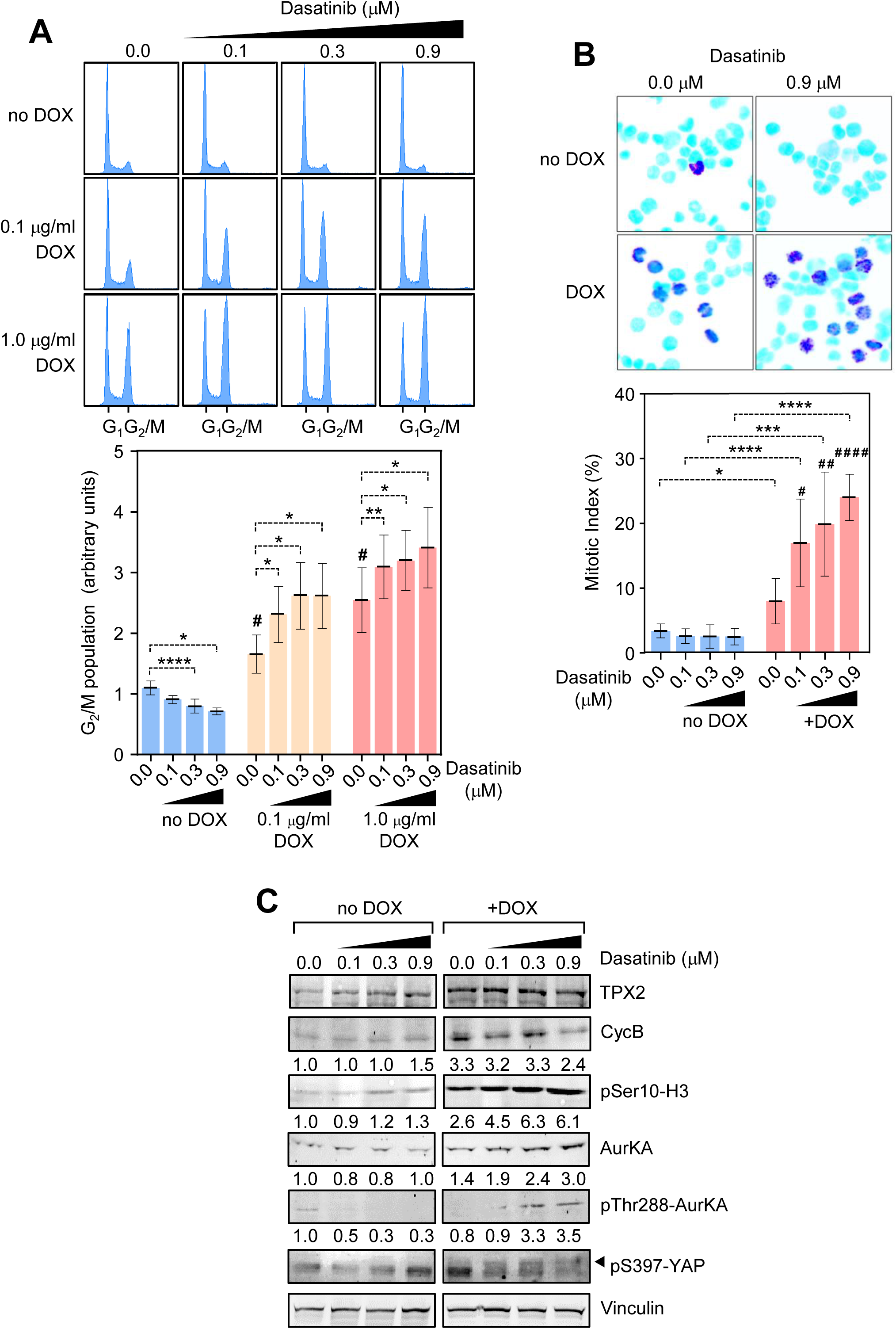
Dasatinib leads to mitotic arrest in TPX2-expressing cells. **A).** Cell cycle profiling by DAPI DNA staining and flow cytometry analysis of MDA-MB-453 cells. Cells incubated with DOX (0.01 μg/ml and 0.1 μg/ml) for TPX2 expression, were also treated with 0.1μM, 0.3μM and 0.9μM of Dasatinib. The G1 and G2/M peaks are indicated at the bottom of the cell cycle profiles, and the quantification of the G2/M percentage of cells is shown in the bottom histogram. Two-way ANOVA with Tuckey multiple comparisons test comparing dasatinib impact: p<0.0001 (****) p<0.01 (**), p<0.05 (*); or TPX2 expression impact versus no DOX treated cells: p<0.05 (#). (blue = no DOX control, orange = 0.1 μg/ml DOX, red = 1.0 μg/ml DOX). **B).** Mitotic index quantification by phospho-Ser10 Histone H3 (pH3) immunofluorescence upon DOX incubation and dasatinib treatment. The upper panel shows a representative image of MDA-MB.453 cells stained with DAPI for DNA (light blue) and pH3 depicting mitotic cells (dark purple). The bottom histogram shows the quantification at different concentrations of dasatinib. Two-way ANOVA with Tuckey multiple comparisons test, comparing TPX2 expression impact: p<0.0001 (****) p<0.001 (***), p<0.05 (*); or dasatinib impact with in the TPX2 expressing cohort; p<0.0001 (####) p<0.01 (##) p<0.05 (#). (blue = no DOX control, orange = 1.0 μg/ml DOX). **C).** Biochemical analysis by western blot of mitotic markers activation, and YAP-Ser397 phosphorylation, of MDA-MB-453 cells expressing TPX2 (DOX) or control cells (ctrl), treated with 0.1μM, 0.3μM, and 0.9μM of Dasatinib. Numbers at the bottom of the WB panels represent normalized band intensity quantified with the LiCor Odyssey 2.0 software. GAPDH expression levels are used as a loading control.

Mitotic arrest can also influence the Hippo-YAP/TAZ signaling in different ways. On one hand, the mitotic kinase CDK1 phosphorylates and activates YAP during the G2/M phase [92, 93]. On the other hand, CDK1 phosphorylates and inactivates TAZ during mitosis [94], and also phosphorylates and activates LATS kinases upon mitotic stress [95, 96]. Other mitotic kinases such as AurKA can modulate the Hippo-YAP/TAZ axis signaling by phosphorylating YAP-Ser397 [97]. Noteworthy, phosphorylation at YAP-Ser397 is controversial in terms of activity, as it plays as a YAP inhibitory event leading to YAP degradation [98], but also as a YAP activator mechanism, precisely upon TPX2 expression and AurKA phosphorylation in breast cancer cell lines [97].

We, therefore, evaluated the levels of YAP-Ser397 phosphorylation, detecting an increased level in YAP-Ser397 phosphoresidue when TPX2 expressing cells are cultured under dasatinib, whereas control cells do not alter YAP-Ser397 phosphorylation levels (Figure 4C). This YAP-Ser397 phosphorylation is coincident with the recovery of the inhibitory YAP-Ser127 signal and reduction of the active YAP signal (Figure 3B), and the cytoplasmic retention (Figure 3C), suggesting that in this context YAP-Ser397 reflects inhibition of YAP activity.

Overall, dasatinib leads to an efficient cell cycle arrest in mitosis in TPX2-expressing cells, leading to YAP inactivation by phosphorylation at Ser397 and Ser127 residues.

### 3.5. TPX2 expression and YAP activation correlation analysis in breast cancer patient-derived samples

To evaluate the clinical translation of our findings, we examined whether TPX2 expression levels and YAP activation (by YAP-pSer127 inhibitory phosphorylation) are coincident biomarkers in an array of 99 invasive ductal breast carcinoma samples (Figure 5A). We first tested that YAP-pSer127 staining inversely correlates with total YAP levels (not shown), thus validating the YAP-pSer127 labeling. Although a fraction of samples show low YAP-pSer127 and no TPX2 expression, the majority (54%) of the TPX2-positive cases also have increased YAP activity (Figure 5B). We also compared other clinical features such as distant metastasis and subtype classification. 70% of patients with distant metastasis have negative YAP-pSer127 staining, indicating that YAP activation is also a hallmark of aggressiveness. Since MDA-MB-453 is a Her2-positive cell line, we also evaluated the HER2 status in the analyzed tumors, showing most cases being negative for YAP-pSer127 staining.

**Figure 5.**
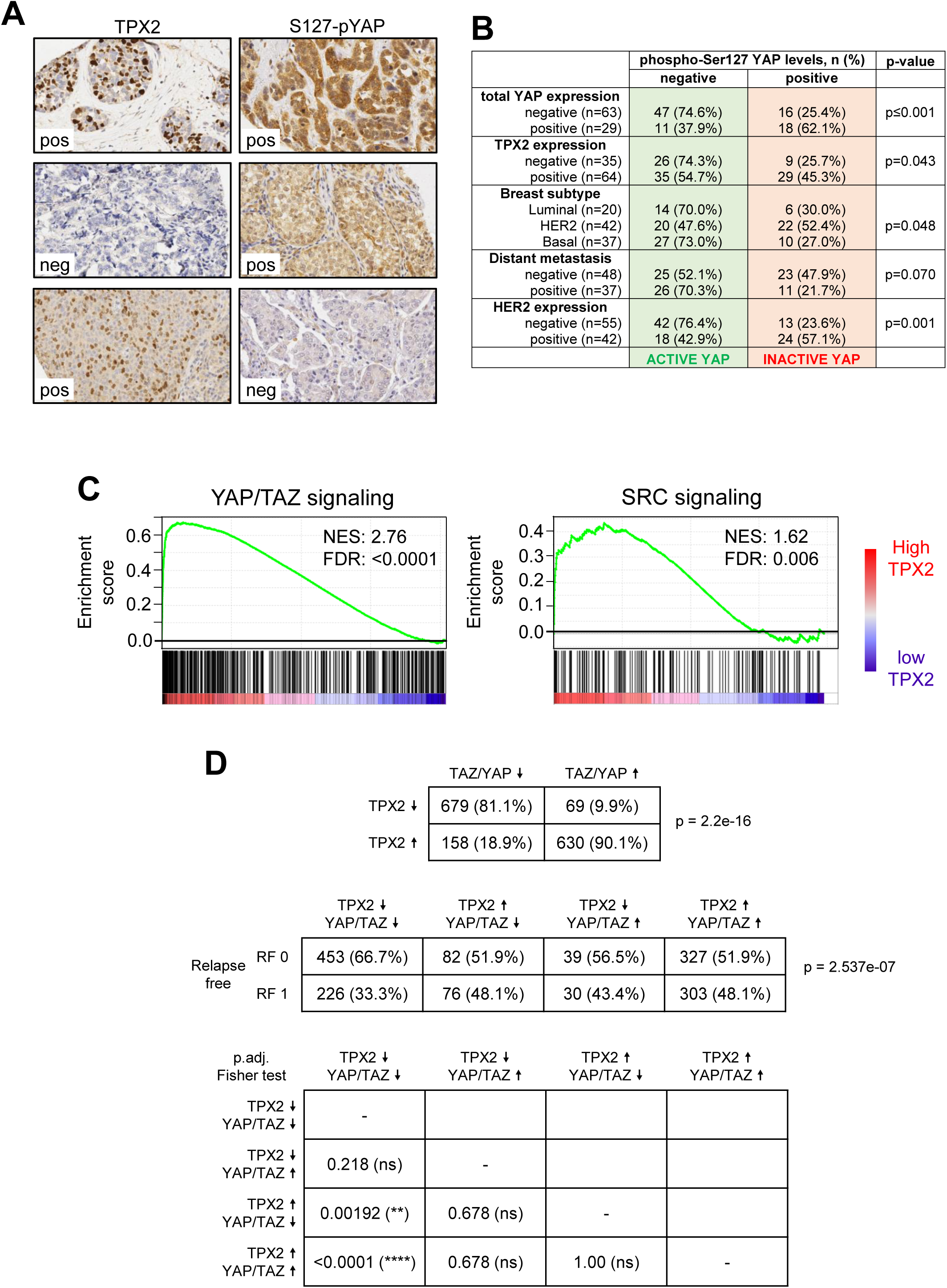
Correlation analysis of TPX2 expression and YAP activation of a breast cancer tumoral microarray. **A).** Immunohistochemistry of TPX2 and pS127-YAP in grade 3 breast cancer paraffin-embedded samples. The upper row shows an example of breast carcinoma with a positive signal for both biomarkers. The middle row is an example of breast carcinoma with no TPX2 expression and with S127-pYAP positive signal (indicative of YAP inactivation). The bottom panel indicates an example of breast carcinoma with positive expression of TPX2 and negative S127-pYAP expression (indicative of YAP activation). Pictures were obtained at 40x magnification. **B).** Quantification and statistical analysis table of 99 samples stained as in panel A showing the relationship between pS127-YAP signal and other immunohistochemical features, such as expression of YAP, TPX2 and HER2, metastasis capacity, and tumoral subtypes. **C).** Gene Set Enrichment Analysis (GSEA) of breast invasive ductal carcinoma samples from the METABRIC project, accordingly to TPX2 expression levels, showing enhanced YAP/TAZ signaling [52] and SRC signaling (SRC_UP.V1_DN) [111]. Normalized Enrichment Score (NES) and FDR qValue are indicated in each plot. **D).** Correlation analysis of TPX2 and the YAP/TAZ signature expression (z-score) in breast cancer samples from the METABRIC project, evaluating the influence on Relapse Free survival (RF0 = no relapse; RF1 = positive relapse). Statistical analysis by adjusted Fisher test: p<0.0001 (****); p<0.01 (**).

To reinforce this data, we explored the METABRIC Project database [50] by performing a Gene Set Enrichment Analysis (GSEA) of 1536 breast invasive ductal carcinoma samples according to the TPX2 expression levels. TPX2 high expression correlates with an increase in YAP/TAZ and SRC-dependent signaling (Figure 5C), and we obtained similar data using the TCGA database (supplementary figure 4B). The METABRIC dataset indicates that the expression of TPX2 and the YAP/TAZ signature strongly correlate (Figure 5D), and has a significant impact on malignancy as depicted by Relapse Free (RF) survival. The few samples where TPX2 and YAP/TAZ signatures do not correlate show no differences in RF, thus suggesting that TPX2 expression needs an elevated YAP/TAZ signaling to provide a poor tumoral prognosis. In summary, breast cancer samples that show high levels of TPX2 often present enhanced YAP activation, making dasatinib a therapeutic option, especially in the more aggressive tumors.

## 4. Discussion

Chromosomal Instability is a hallmark of cancer, being a feature not present in healthy primary cells. Therefore, it might be a bona fide biomarker for cancer therapy, and strong efforts have been done trying to find drugs that selectively kill cancer cells based on their CIN rate [8, 9, 99]. Despite some of these studies show positive correlation between CIN and certain therapies [99], the final results generally demonstrated that CIN is difficult to target, and the few drugs depicted are very toxic compounds [9]. In addition, high-CIN cancer cells often show elevated proliferation rates, and this feature can be responsible for positive responses to drugs based on protein turnover or energy balance [8]. The fact that CIN cells are already very heterogeneous in their genetic dosage can explain why they do not respond to therapy uniquely based on the CIN rate. Moreover, this genetic plasticity provides adaptation to external insults, being an engine for therapy resistance [100, 101].

CIN is characterized by the expression of particular genetic signatures, mainly based on mitotic and cell cycle genes [10–16]. Using a gene-inducible expression system, we screened cancer cells, individually expressing a collection of CIN-related genes, against a selected library of drugs. This approach avoids the enormous differences between different cell lines, as we compare the same cell line expressing or not the gene of interest. Thus, the genetic background is the same, and we can depict the precise therapy dependency on one gene. We found that the expression of TPX2, a major representative gene in CIN signatures, provides strong sensitivity towards SRC kinase inhibitors such as dasatinib, saracatinib, and bosutinib (Figure 1). Interestingly, none of the other tested genes provided such a response. This indicates that the drug response is TPX2 specific rather than CIN associated. Indeed, there is no significant correlation between dasatinib response and CIN rate, or expression of other CIN-related genes, according to the DepMap database (Supplementary Figure 2). Worth mentioning, other research groups have shown that SRC inhibitors can affect the proliferation of CIN cells, although not as a general mechanism. Firstly, Schukken and coworkers made a similar screen approach by confronting a small panel of drugs in cells depleted for MAD2 (leading to severe aneuploidy and CIN in the short term) and showed an increased sensitivity towards the SRC inhibitor SKI606 (bosutinib) [102]. Here, bosutinib exacerbates the mitotic aberrations generated upon MAD2 depletion, by increasing the polymerization rates of microtubules, evidencing similarities to our cell cycle analysis showing how dasatinib leads to a strong mitotic arrest upon TPX2 expression (Figure 4). Whether this alteration depends on microtubule dynamics, in the TPX2-expressing cells, remains to be determined. Secondly, in an exhaustive in silico study using single-nucleotide polymorphism (SNP) data from the TCGA, researchers identified 17 different CIN signatures aiming to predict drug response and new drug targets [103]. Interestingly, this study describes that dasatinib precisely correlates with a CIN signature (CX10), which originated from a defective non-homologous end joining (NHEJ) DNA repairing mechanism. The TPX2/AurKA complex is implicated in DNA repair, as it interacts with essential DNA repair factors such as BRCA1/2, PARP1, and 53BP1. TPX2 is known to accumulate at DSBs where it negatively regulates 53BP1, thus modulating homologous recombination (HR), replication fork stability, and inhibiting NHEJ [104–107].

The association between SRC and the Hippo-YAP pathway is well documented in the literature [43], and a tightly controlled SRC-YAP signaling axis determines therapeutic response to dasatinib mediated by the stress kinase JNK [45]. Since we also observed that the stress kinase JNK is activated upon TPX2 expression and dasatinib addition leads to a significant reduction of JNK activation (Figure 2), we deeply analyzed the impact of TPX2 expression on YAP signaling, demonstrating that TPX2 high levels promote YAP activation and this is probably the reason for dasatinib-increased sensitivity (Figure 3). Our data is concomitant with previous works showing that TPX2 expression increases YAP signaling in breast cancer cells, and this is dependent on Aurora A kinase activity [81, 82, 97]. Moreover, Aurora A is also known for modulating SRC through the cofactor NEDD9, affecting dasatinib response [108].

The reduced proliferation in TPX2-expressing cells, upon dasatinib treatment, can be explained by the strong mitotic arrest observed (Figure 4). Although SRC kinases are primarily implicated in mitogenic signaling, there is also evidence that they can modulate mitosis. SRC-family kinases modulate microtubule polymerization rates [102, 109], a process where TPX2 also has an important role. Similarly, v-SRC expression overrides the spindle assembly checkpoint leading to chromosome missegregation [91, 110]. The mitotic alterations generated by TPX2 overexpression can synergize with the inhibition of SRC by dasatinib, leading to the observed mitotic arrest, and further studies need to be done to determine the precise mechanism of action.

Intending to translate our data to real cancer samples, we explored the breast cancer data from the METABRIC project, and a cohort of breast invasive ductal carcinoma samples (Figure 5). The data obtained show that indeed elevated TPX2 strongly correlates with increased SRC and YAP signaling and that TPX2 confers poor prognosis only when YAP signaling is elevated. About 54% of TPX2-positive breast tumors also harbor YAP activation, demonstrating that both biomarkers are present in a significant proportion of breast tumors. Interestingly, these tumors seem to be the most aggressive. Collectively, our data suggest TPX2 expression and YAP signaling as putative biomarkers for alternative dasatinib therapy in aggressive breast cancer.

## 5. Conclusions

- TPX2 overexpression provides enhanced sensitivity to the SRC kinase inhibitor dasatinib.
- This increased dasatinib sensitivity is based on YAP/TAZ signaling activation upon TPX2 expression.
- Dasatinib leads to an efficient mitotic arrest in TPX2-expressing cells
- Analysis in patient-derived breast cancer samples shows a strong correlation between high TPX2 expression levels and YAP/TAZ signaling activation, and this provides a poor prognosis.

## 6. Acknowledgements

We thank Prof. Erch Nigg for sharing the HEC1 antibody. This study has been supported by the following grants: From the “Ministerio de Ciencia, Innovación, Agencia Estatal de Investigación MCIN/AEI/FEDER (doi:10.13039/501100011033): RTI2018-095496-B-I00 and PID2021-125705OB-I00 (GdC); PID2019-104644RB-I00 & PID2022-136854OB-I00 (GMB); Juan de la Cierva Postdoctoral program FJC2020-044620-I (NSG). From the Spanish Association Against Cancer (AECC) Scientific Foundation: LABAE16017DECA (GdC); PROYE19036MOR (GMB). From the Spanish National Research Council (CSIC): 2018-20I114, 2021-AEP035, and 2022-20I018 (GdC). From the Instituto de Salud Carlos III – CIBERONC: CB16/12/00295 (GMB).

## 7. Author contributions

CM performed the drug screening and the in vitro experiments, with the help of BO, AMV, CB, MG, and AAdlV. AT and GMB performed the breast tumoral samples analysis. NSG performed the METABRIC and TCGA bioinformatic analysis. MJL provided intellectual input and supervised the drug screening. GdC designed and supervised the study. All authors participated in the data analysis, and GdC wrote the paper with the help of NSG, CM, and MJL

## 8. Conflict of interests

Carlos Marugán and Maria José Lallena are employees and shareholders of Eli Lilly and Company.

## Notes

### Competing Interest Statement

Carlos Marugan and Maria Jose Lallena are employees and shareholders of Eli Lilly and Company.

