## Supplementary Figures and Tables for "TPX2 expression promotes sensitivity to dasatinib in breast cancer by activating the YAP transcriptional signaling"

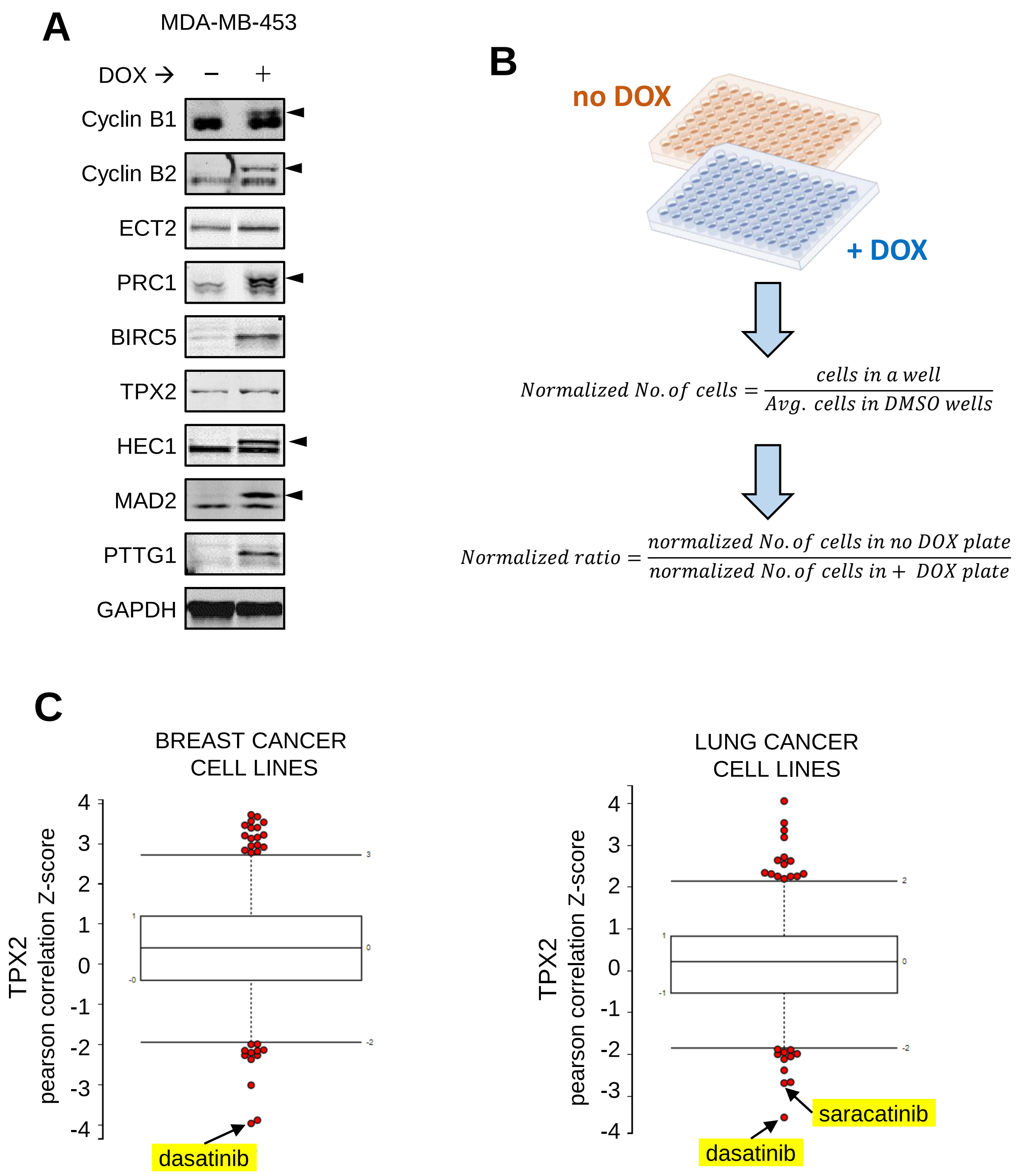
A
MDA-MB-453
DOX →
−
+
Cyclin B1
Cyclin B2
ECT2
PRC1
BIRC5
TPX2
HEC1
MAD2
PTTG1
GAPDH
B
no DOX
+ DOX
$Normalized\ No.of\ cells = \frac{cells\ in\ a\ well}{Avg.\ cells\ in\ DMSO\ wells}$
$Normalized\ ratio = \frac{normalized\ No.of\ cells\ in\ no\ DOX\ plate}{normalized\ No.of\ cells\ in +\ DOX\ plate}$
C
BREAST CANCER CELL LINES
TPX2 pearson correlation Z-score
4
3
2
1
0
-1
-2
-3
-4
dasatinib
LUNG CANCER CELL LINES
TPX2 pearson correlation Z-score
saracatinib
dasatinib

Supplementary Figure 1

**A)** Inducible expression test for the nine cDNAs in MDA-MB-453 cells upon doxycycline addition (1.0μg/ml) for 24 hours.

**B)** Cartoon showing how the drug screen was performed in MDA-MB-453 cells. 10 serial dilutions of each drug are added to cells in duplicates, and plates are treated either with doxycycline media or with regular media for cDNA expression. Drug incubation lasted two doubling times, and DNA stained cells were counted. The number of cells was normalized by the average cell number in the DMSO-treated cells (no drug), and then the ratio of no DOX/+ DOX was calculated for each drug concentration.

**C)** Z-scored Pearson correlation coefficients between small-molecule sensitivity data, expressed as areas under concentration-response curves (AUCs), with basal TPX2 gene-expression measurements, expressed as log2 robust-multi-array-average values, in Breast Cancer and Lung Cancer cell lines, retrieved from the Cancer Therapeutics Response Portal (CTRP). (https://portals.broadinstitute.org/ctrp.v2.1/?featureName=TPX2)

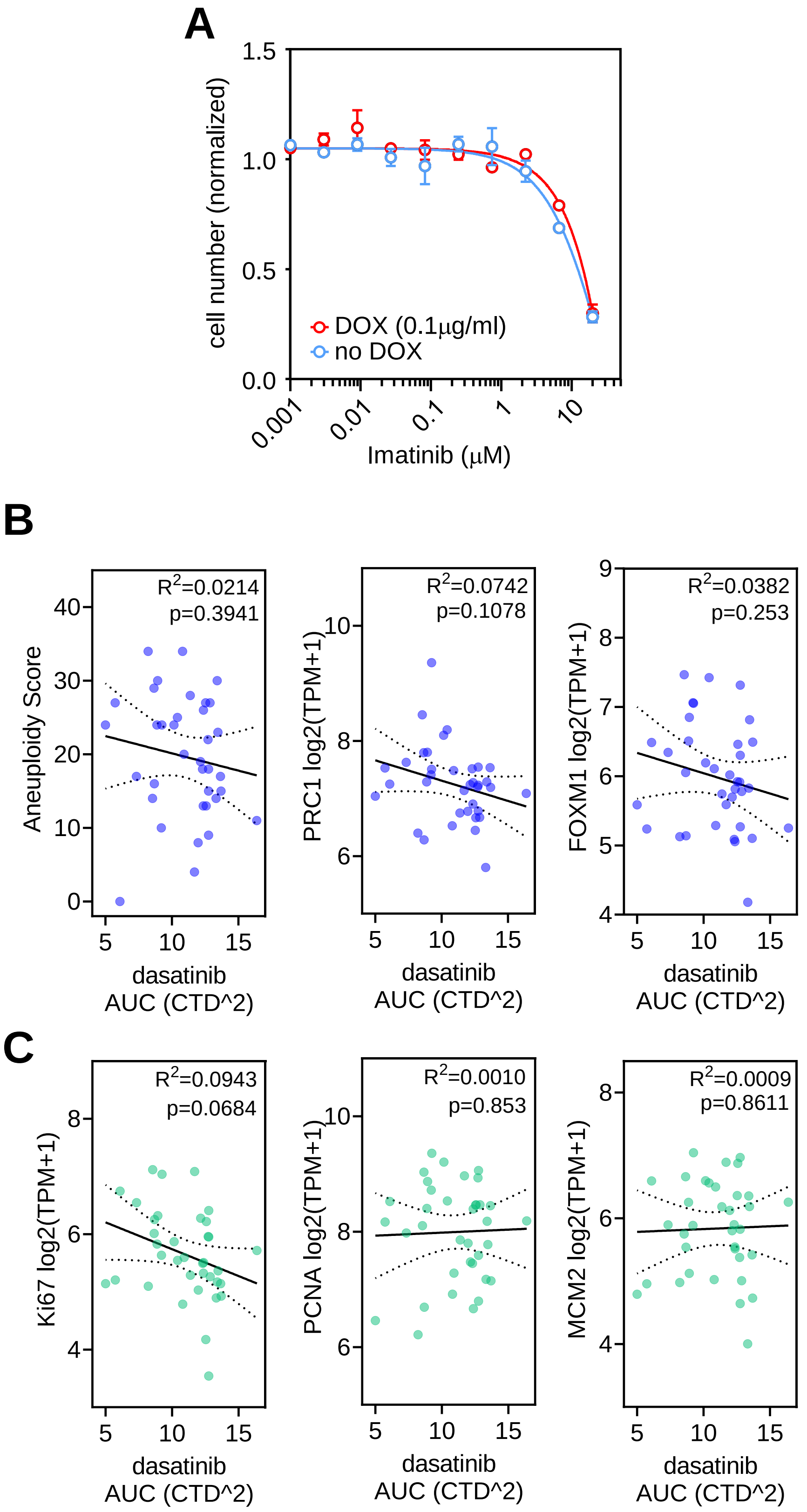
A
cell number (normalized)
1.5
1.0
0.5
0.0
DOX (0.1µg/ml)
no DOX
0.001
0.01
0.1
1
10
Imatinib (µM)
B
Aneuploidy Score
R2=0.0214
p=0.3941
PRC1 log2(TPM+1)
R2=0.0742
p=0.1078
FOXM1 log2(TPM+1)
R2=0.0382
p=0.253
dasatinib
AUC (CTD^2)
C
Ki67 log2(TPM+1)
R2=0.0943
p=0.0684
PCNA log2(TPM+1)
R2=0.0010
p=0.853
MCM2 log2(TPM+1)
R2=0.0009
p=0.8611
dasatinib
AUC (CTD^2)

Supplementary Figure 2

**A)** 10-point Concentration-Response Curves (CRC) and IC50 calculation of imatinib, in MDA-MB-453 cells expressing TPX2, during 3 doubling times. Mean cell number normalized vs. DMSO +/- SEM (blue = no DOX control, red = 0.1 μg/ml DOX).

**B)** DepMap portal retrieved data from breast cancer cell lines, showing the correlation between the sensitivity to dasatinib (area under the curve - AUC) versus the aneuploidy score (based on the ABSOLUTE copy number data from the Cancer Cell Line Encyclopedia (*Ghandi, M., et al. Nature 2019. v569, 503–508. doi: 10.1038/s41586-019-1186-3*)), or the expression of the CIN related genes PRC1 and FOXM1 in transcripts per million (TPM).

**C)** Similar DepMap portal correlation analysis of dasatinib sensitivity (AUC) versus the expression levels (TPM) of the proliferating genes MKI67, PCNA or MCM2

Suppl. Figure 3 Marugan et al.

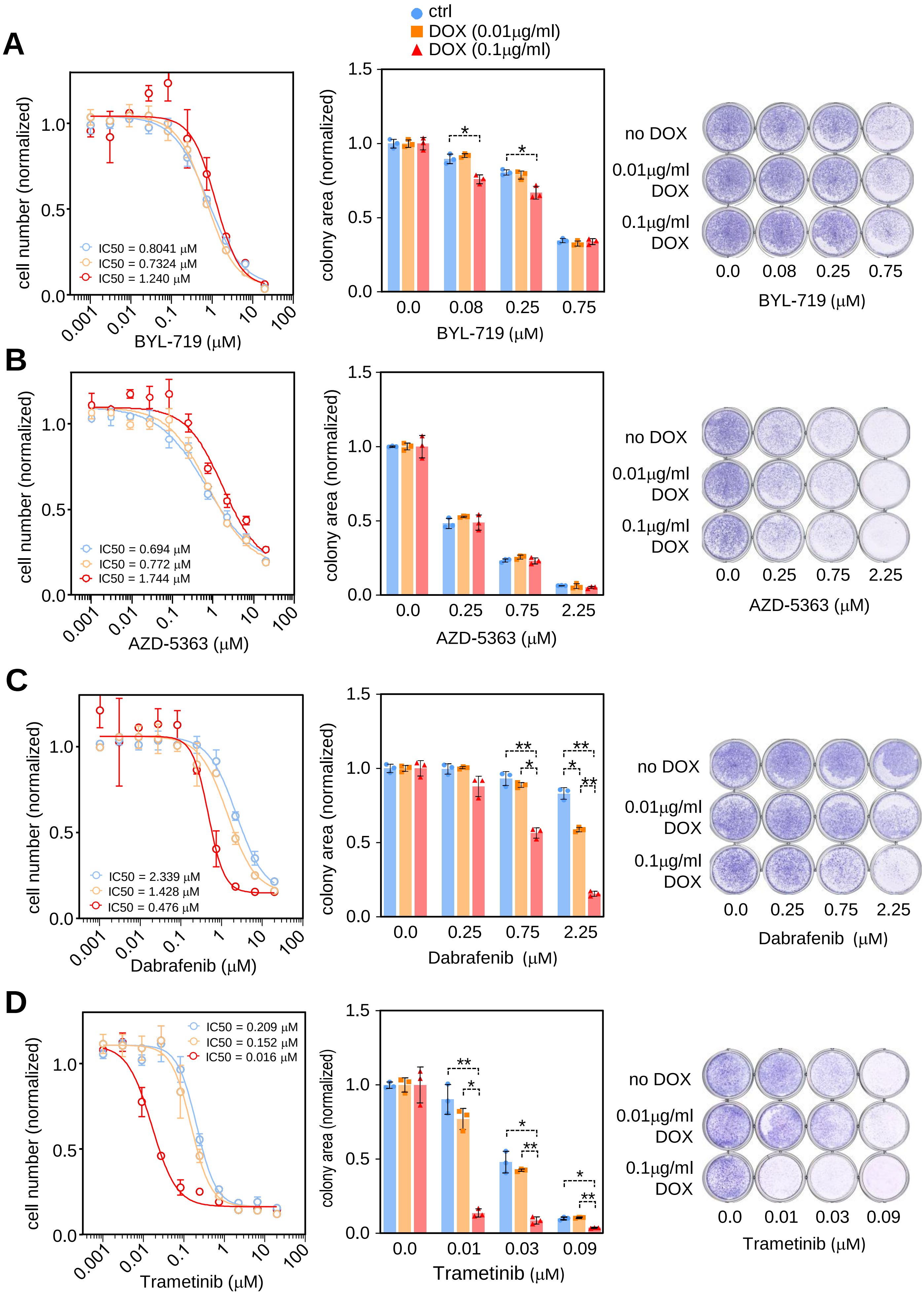

Supplementary Figure 3

Response of TPX2-expressing cells to PI3K/AKT and RAS/MEK/ER inhibitors.

**A)** PI3K kinase inhibitor BYL-719.

**B)** AKT inhibitor AZD-5363.

**C)** RAF inhibitor dabrafenib.

**D)** MEK inhibitor trametinib.

**Left Panels:** 10-point CRC and IC50 calculation of the indicated drug in MDA-MB-453 cells expressing TPX2, during 3 doubling times. Mean cell number normalized vs. DMSO +/- SEM (blue = no DOX control, orange = 0.01 μg/ml DOX, red = 0.1 μg/ml DOX).
**Middle panels:** Colony formation assay, during two weeks in MDA-MB-453 cells upon treatment with the indicated inhibitors. The colony area is normalized vs. the DMSO-treated cells +/- SD. Two-way ANOVA with Tuckey multiple comparisons test: $p<0.01$ (**), $p<0.05$ (*). (blue = no DOX control, orange = 0.01 μg/ml DOX, red = 0.1 μg/ml DOX).
**Right panels:** representative image of the colony formation assay.

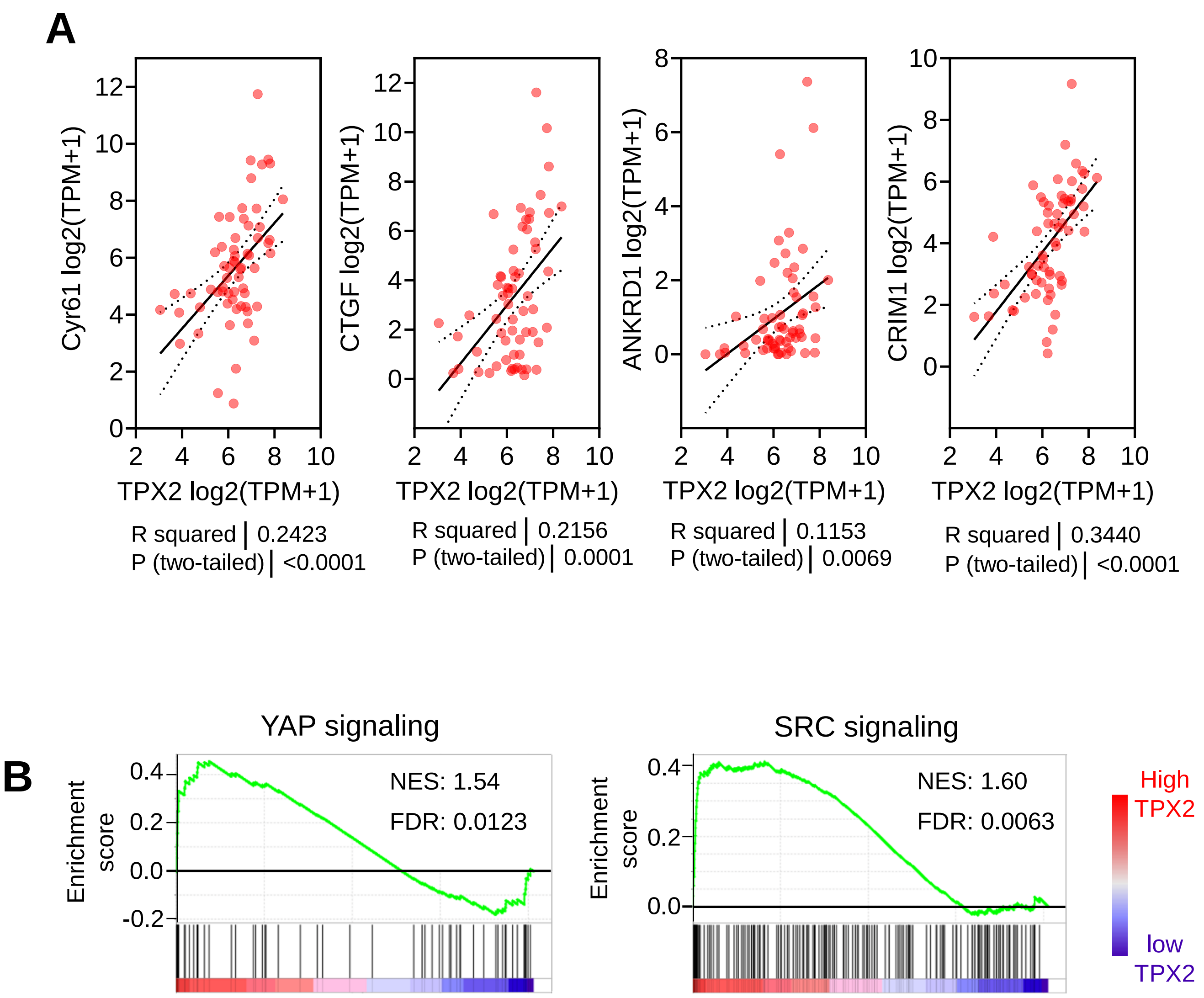
A
Cyr61 log2(TPM+1)
CTGF log2(TPM+1)
ANKRD1 log2(TPM+1)
CRIM1 log2(TPM+1)
TPX2 log2(TPM+1)
R squared | 0.2423
P (two-tailed) | <0.0001
R squared | 0.2156
P (two-tailed) | 0.0001
R squared | 0.1153
P (two-tailed) | 0.0069
R squared | 0.3440
P (two-tailed) | <0.0001
B
YAP signaling
SRC signaling
Enrichment score
NES: 1.54
FDR: 0.0123
NES: 1.60
FDR: 0.0063
High TPX2
low TPX2

Supplementary Figure 4

**A).** Correlation analysis of TPX2 and YAP/TAZ signaling surrogate genes (Cyr61, CTGF, ANKRD1, and CRIM1) expression levels in transcripts per million (TPM), using breast cancer cell lines data retrieved from the DepMap portal.

**B).** Gene Set Enrichment Analysis (GSEA) of 1082 patients breast invasive ductal carcinoma samples from the TCGA-PanCanAtlas and BRCA project (*Cancer Genome Atlas Research. Nat Genet. 2013. v45(10):1113-20. doi: 10.1038/ng.2764*), accordingly to TPX2 expression levels, showing enhanced YAP/TAZ signaling. (Cordenonsi_YAP_Conserved_Signature) (*Cordenonsi et al., Cell. 2011 v147(4):759-72. doi: 10.1016/j.cell.2011.09.048)* and SRC signaling (SRC_UP.V1_DN) (*Bild. A.H, et al., Nature. 2006. v439(7074):353-7. doi: 10.1038/nature04296*). Patients were classified according to TPX2 expression with a membership probability estimated by bootstrap (*Bueno-Fortes S, et al., Bioinform Adv. 2023. v3(1). doi: 10.1093/bioadv/vbad037*). Gene Set Enrichment Analysis (GSEA) was performed by comparing high versus low TPX2 expressing samples. The top 25 terms of C6: Oncogenic Signature were analyzed. FDR q< 0.05 was considered significant.

**Supp. Table 1. Screening drug collection.**

The table summarizes the collection of 60 small compounds used in the screening, by name, target molecule, and the clinical status (cpd: compound).

| # | Inhibitor | Target | Status |
|---|---|---|---|
| 1 | Afatinib | EGFR, HER2 | FDA approval |
| 2 | Axitinib | c-Kit, VEGFR, PDGFR | FDA approval |
| 3 | Bosutinib | SRC | FDA approval |
| 4 | Crizotinib | c-Met, ALK | FDA approval |
| 5 | Dabrafenib | Raf | FDA approval |
| 6 | Dasatinib | BCR-ABL, SRC | FDA approval |
| 7 | Erlotinib | Autophagy, EGFR | FDA approval |
| 8 | Everolimus | mTOR | FDA approval |
| 9 | Gefitinib | EGFR | FDA approval |
| 10 | Ibrutinib | BTK | FDA approval |
| 11 | Imatinib | PDGFR | FDA approval |
| 12 | Lapatinib | HER2, EGFR | FDA approval |
| 13 | Palbociclib | CDK4 | FDA approval |
| 14 | Rapamycin | Autophagy, mTOR | FDA approval |
| 15 | Ribociclib | CDK4 | FDA approval |
| 16 | Ruxolitinib | JAK | FDA approval |
| 17 | Sorafenib | Raf | FDA approval |
| 18 | Tofacitinib | JAK | FDA approval |
| 19 | Trametinib | MEK | FDA approval |
| 20 | Vemurafenib | Raf | FDA approval |
| 21 | AZD-1208 | Pim | Phase I |
| 22 | AZD-7762 | Chk | Phase I |
| 23 | JNJ-38877605 | c-Met | Phase I |
| 24 | MK-5108 | Aurora | Phase I |
| 25 | Mubritinib | HER2 | Phase I |
| 26 | PF-00562271 | FAK | Phase I |
| 27 | YM155 | BIRC5 | Phase I |
| 28 | Acadesine | AMPK | Phase II |
| 29 | Apatinib | VEGFR | Phase II |
| 30 | AZD-5363 | Akt | Phase II |
| 31 | BYL-719 | PI3K | Phase II |
| 32 | Flavopiridol | CDKs (CDK1/2/4/6/7) | Phase II |
| 33 | Refametinib | MEK | Phase II |
| 34 | Roscovitine | CDKs (CDK1/2/5) | Phase II |
| 35 | Sotrastaurin | PKC | Phase II |
| 36 | Neflamapimod | p38 MAPK | Phase II |
| 37 | AZD-4547 | FGFR | Phase II/III |
| 38 | Barasertib | Aurora | Phase II/III |
| 39 | Alisertib | Aurora | Phase III |
| 40 | Bardoxolone Methyl | IκB/IKK | Phase III |
| 41 | Crenolanib | PDGFR | Phase III |
| 42 | Dacomitinib | EGFR | Phase III |
| 43 | Dinaciclib | CDKs (CDK2/5/1/9) | Phase III |
| 44 | Dovitinib | FGFR, FLT3, c-Kit, VEGFR, PDGFR | Phase III |

| | | | |
|---|---|---|---|
| 45 | Ceritinib | ALK | Phase III |
| 46 | Linsitinib | IGF-1R | Phase III |
| 47 | Losmapimod | p38 MAPK | Phase III |
| 48 | Quizartinib | FLT3 | Phase III |
| 49 | Volasertib | PLK1, PLK2, PLK3 | Phase III |
| 50 | 10058-F4 | c-Myc | preclinical |
| 51 | 5-Iodotubercidin | Haspin, ADK | preclinical |
| 52 | AR-A014418 | GSK-3 | preclinical |
| 53 | EHT-1864 | Rho | preclinical |
| 54 | GNE-7915 | LRRK2 | preclinical |
| 55 | GSK429286A | ROCK | preclinical |
| 56 | KU-60019 | ATM/ATR | preclinical |
| 57 | Nocodazole | Microtubules | reference cpd |
| 58 | Vinblastine | Microtubules | reference cpd |
| 59 | Paclitaxel | Microtubules | reference cpd |
| 60 | Staurosporine | multiple kinases | reference cpd |

**Supp. Table 2. cDNA cloning oligos**

The table shows the gene name, accession number, species of origin, and the forward (FW) and reward (RW) used oligos for PCR. The red CACC indicates the pENTR/D-topo orientation cloning sequence.

| Gene | Access Number | Species | 5'-sequence-3' | |
|---|---|---|---|---|
| BIRC5 | BC008718 | human | FW | CACCATGGGTGCCCCGACGT |
| | | | RW | ATCCATGGCAGCCAGCTG |
| *Ccnb1* | NM_172301.3 | mouse | FW | CACCATGGCGCTCAGGGTCAC |
| | | | RW | TGCCTTTGTCACGGCCTTAG |
| *Ccnb2* | BC008247 | mouse | FW | CACCATGGCGCTGCTCCGAC |
| | | | RW | GGGGCTGCCCAGCA |
| ECT2 | BC112086 | human | FW | CACCATGGCTGAAAATAGTGTATT |
| | | | RW | TATCAAATGAGTTGTAGATCTAC |
| HEC1 | BC035617 | human | FW | CACCATGAAGCGCAGTTCAGTTTCCA |
| | | | RW | TTCTTCAGAAGACTTAATTAGAGTAG |
| MAD2 | BC000356 | human | FW | CACCATGGCGCTGCAGCTCTC |
| | | | RW | GTCATTGACAGGAATTTTGTAGGC |
| PRC1 | BC003138 | human | FW | CACCATGAGGAGAAGTGAGGTGCTG |
| | | | RW | GGACTGGATGTTGGTTGAATTGAG |
| *Pttg1* | BC023324 | mouse | FW | CACCATGGCTACTCTTATCTTTGTTGA |
| | | | RW | AATATCTGCATCGTAACAAACAGGTG |
| *Tpx2* | BC060619 | mouse | FW | CACCATGTCACAAGTCCCTACTACTTA |
| | | | RW | CTACTGGAACCGAGTGGAGAACT |

**Supp. Table 3. Antibodies for WB**

Table summarizing all the antibodies used for western blot detection, indicating the protein name, the vendor company and reference, the animal source and clonal type, and dilution used. (SCBT - Santa Cruz Biotechnology; CST - Cell Signaling; CNIO – Spanish National Cancer Center; BD – Becton Dickinson)

| **Protein** | **Vendor** | **Reference** | **Host** | **Dilution** |
|---|---|---|---|---|
| ABL1 | SCBT | sc-131 | Rabbit polyclonal | 1:500 |
| AKT | CST | #9272 | Rabbit polyclonal | 1:500 |
| AKT-pSer473 | CST | #4060 | Rabbit polyclonal | 1:1000 |
| AurKA | Abcam | ab13824 | Mouse monoclonal | 1:500 |
| AurKA-pThr288 | CST | #3079 | Rabbit monoclonal | 1:500 |
| BIRC5 | Novus | NB 500-201 | Rabbit polyclonal | 1:1000 |
| Cyclin B1 | SCBT | sc-752 | Rabbit polyclonal | 1:1000 |
| Cyclin B2 | Abcam | ab185622 | Rabbit monoclonal | 1:1000 |
| ECT2 | SCBT | sc-1005 | Rabbit polyclonal | 1:500 |
| ERK | CST | #9102 | Rabbit polyclonal | 1:1000 |
| ERK-pThr202/204 | CST | #9101 | Rabbit polyclonal | 1:1000 |
| GAPDH | CNIO | FF26A | Mouse monoclonal | 1:5000 |
| HEC1 | Erich Nigg lab | | Rabbit polyclonal | 1:2000 |
| Histone H3-pSer10 | CST | #3377S | Rabbit monoclonal | 1:1000 |
| JNK-pThr183/pTyr185 | CST | #9251 | Rabbit polyclonal | 1:500 |
| LATS1 | SCBT | sc-398560 | Mouse monoclonal | 1:500 |
| LATS1-pSer909 | CST | #9157 | Rabbit polyclonal | 1:500 |
| MAD2 | BD | 610679 | Mouse monoclonal | 1:500 |
| p38 | SCBT | sc-7972 | Mouse monoclonal | 1:500 |
| p38-pTyr182 | SCBT | sc-166182 | Mouse monoclonal | 1:500 |
| PRC1 | Abcam | ab51248 | Rabbit monoclonal | 1:1000 |
| PTTG1 | Abcam | ab3305 | Mouse monoclonal | 1:1000 |
| SRC | SCBT | sc-8056 | Mouse monoclonal | 1:500 |
| SRC-pTyr416 | CST | #6943 | Rabbit monoclonal | 1:1000 |
| STAT3-pTyr705 | SCBT | sc-8059 | Rabbit monoclonal | 1:500 |
| TAZ | CST | #4883S | Rabbit polyclonal | 1:1000 |
| TPX2 | Abcam | ab32795 | Mouse monoclonal | 1:500 |
| Vinculin | SCBT | sc-73614 | Mouse monoclonal | 1:2000 |
| YAP | Abcam | ab62751 | Rabbit polyclonal | 1:500 |
| YAP-pSer127 | CST | #4911S | Rabbit polyclonal | 1:1000 |
| YAP-pSer397 | CST | #13619 | Rabbit monoclonal | 1:500 |
